# The maintenance and de-mixing of extrachromosomal DNA variants in single cells

**DOI:** 10.1101/2024.10.22.619675

**Authors:** Elisa Scanu, Benjamin Werner, Weini Huang

## Abstract

Extrachromosomal DNA (ecDNA) has emerged as a key driver of oncogene amplification and a major contributor to rapid intra-tumour heterogeneity, thereby promoting tumour progression and therapeutic resistance. This heterogeneity arises from pronounced cell-to-cell variability in ecDNA copy number, enabling complex ecDNA amplicon compositions within individual tumour cells. Approximately one-third of ecDNA-positive tumours harbour multiple co-selected ecDNA species. However, the mechanisms governing the heterogeneity and persistence of ecDNA variants — beyond the presence of distinct ecDNA species — remain less well understood. In particular, little is known about the maintenance of genetic or phenotypic diversity within a single ecDNA species. Here, we develop computational models to investigate the dynamics that enable the stable maintenance of tumour cells carrying multiple ecDNA variants (“mixed cells”). We explore how variant switching contributes to the persistence of ecDNA diversity under varying fitness regimes. Our results demonstrate that both a positive fitness of ecDNA+ cells and variant switching are required to maintain mixed cell subpopulations, whereas direct co-selection of mixed cells is not necessary. Notably, the fraction of mixed cells peaks at intermediate switching rates across fitness landscapes, a pattern reflected in subpopulation structures, transition probabilities between pure and mixed ecDNA states, and single-cell Shannon diversity indices.

## Introduction

Extrachromosomal DNA (ecDNA) is highly prevalent in human cancers [1]. Although ecDNA has been described for decades in various organisms [2], including human cancers [3–5], the driving role of ecDNA in cancer promotion and progression only became evident recently [6]. ecDNA-positive tumors are among the most aggressive types, are enriched for metastatic disease, and have overall worse clinical outcomes [7, 8].

In part, these observations are explained by the unique evolutionary properties of ecDNAs. ecDNAs segregate randomly between daughter cells [8–11], enabling high cell-to-cell ecDNA copy number variation, high oncogene amplification and rapid evolution [12–14]. Even worse, about 30% of ecDNA positive tumours contain two or more ecDNA species carrying different oncogenes or enhancers [15– 17]. We have previously shown that co-selection of cells containing multi-species ecDNA is necessary to maintain their coexistence [16]. This underscores the complexity of oncogenic processes within cancer cells and facilitates robust inter-ecDNA interactions. For instance, regulatory interactions between ecDNAs, such as enhancer-mediated gene activation, can dramatically increase oncogene expression beyond those achievable by single ecDNA species [15, 17, 18].

It is then natural to extent the concept of ecDNA diversity beyond ecDNA species carrying distinct oncogenes. Identical single-species ecDNAs can acquire point mutations, structural variations and/or epigenetic variations, inducing ecDNA heterogeneity in single cells. The sizes of selected ecDNA amplicons are typically between 1 to 3 Mbp [19]. With a mutation rate of ≈10^−8^ per bp per division, this would imply point mutations of the order of ≈10^−2^ per ecDNA copy per division [20, 21]. Previous studies have shown that mutation rates can be even higher because the repair efficiency is often lower on ecDNA compared to chromosomal DNA, resulting in point mutations and small indels upon DNA damage repair [22, 23]. Some examples are clustered somatic mutations and kyklonic hypermutation, caused by APOBEC3 activity, affecting more than 30% of ecDNA driven cancers [24].

ecDNAs also have higher chromatin accessibility than linear DNA [9, 19, 25, 26], and therefore are highly sensitive to histone modification, such as methylation and acetylation. These are epigenetic mechanisms that can alter chromatine’s structure and genetic expression without modifying the genetic composition of ecDNAs. For example, de-methylation, along with copy number amplification, can increase the expression level of certain genes on ecDNA, such as EGFR [14, 27, 28]. In contrast, methylation can suppress some genes targeted by the human immune system, through for example the KDM5B histone [29], enhancing the epigenetic adaptation of cancer cells.

In a population where ecDNA is randomly segregated, the expected long-term outcome is population de-mixing. That is, depending on the selection regime, the population tends to accumulate in either single-type ecDNA states or becomes ecDNA-free [8, 16]. In this baseline setting, single cell ecDNA heterogeneity is expected to vanish over time. Nonetheless, in the case of ecDNA species we have shown that co-selection enables stable co-existence [16, 21]. Additionally, a recent study has proposed hitchhiking as a mechanism to sustain variation on ecDNAs, though just on intermediate time scales [30].

There is currently no evidence for co-selection in the context of genotypic or phenotypic ecDNA heterogeneity, where differences arise on the same ecDNA rather than different ecDNA species. In such cases, the long-term coexistence of ecDNA variants within a single cell is uncertain. We thus developed a general computational framework and ask when and how this ecDNA heterogeneity can be maintained in single cells, or if mixed cell states driven by random segregation always extinct in the absence of co-selection.

## Methods

### Switching, selection and segregation

We consider the dynamics of two ecDNA variants, represented for simplicity by the colors yellow and red throughout the manuscript (Figure 1a) [21]. We introduce switching rates *p*_*y*_ and *p*_*r*_ in the range [0, 1] that allow ecDNA variants to turn from yellow to red, or red to yellow respectively (Figure 1a). Different biological processes may be captured by different regimes of the switching rates, e.g. the rate to acquire a point mutation per cell division is of the order of 10^−3^ or 10^−2^, while the rate of a phenotypic change may be of the order of 10^−2^ or 10^−1^ per ecDNA per division [14, 31]. ecDNA species correspond to the limiting case of switching rates equal to 0 [16]. Although switching rates could differ for different ecDNA variants in principal, here we focus on the symmetric case where *p*_*y*_ = *p*_*r*_ = *p*.

**Figure 1.**
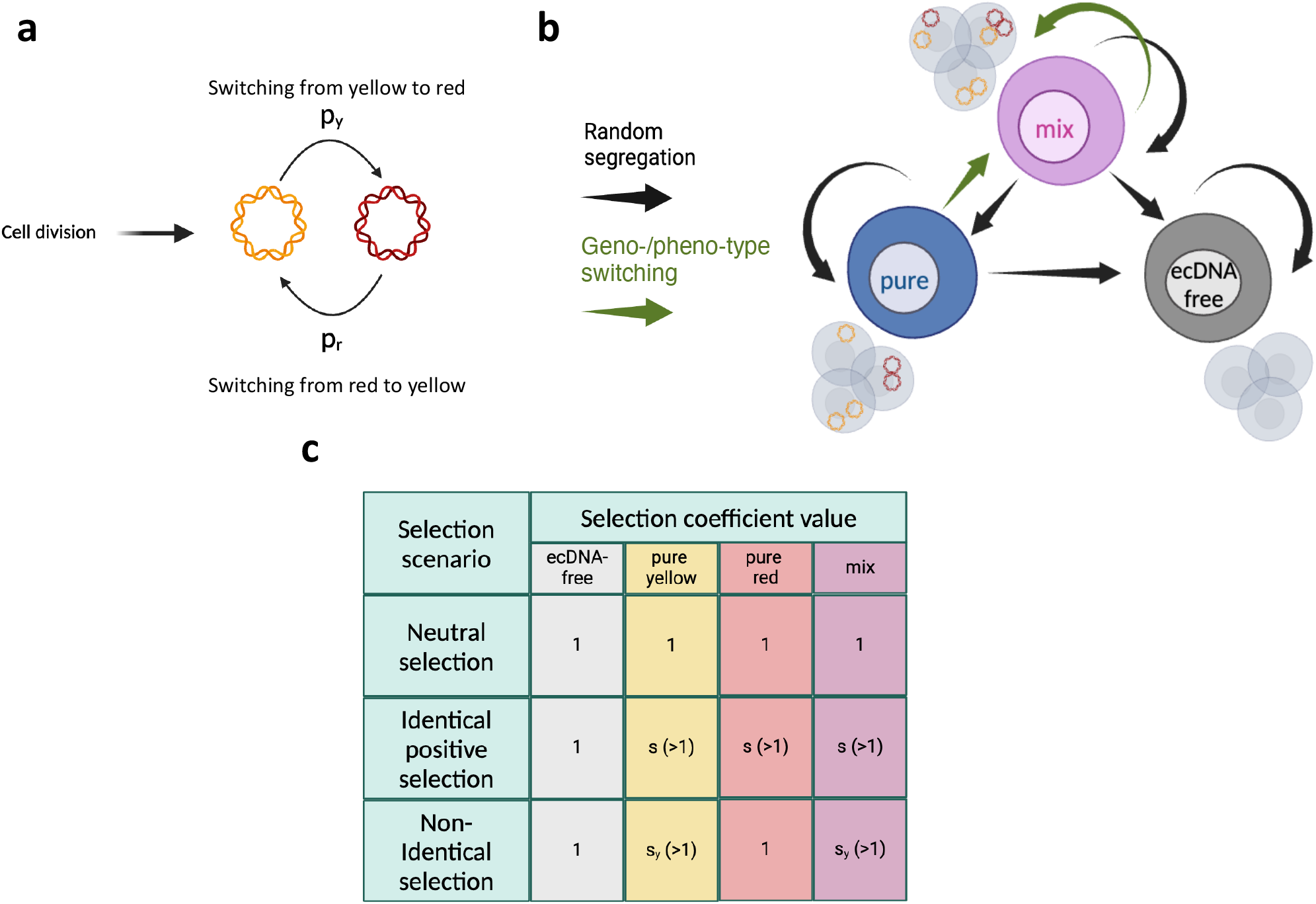
Switching and selection. **a. Switching**. At each cell division, each ecDNA can change colour with probabilities *p*_*y*_ and *p*_*r*_. **b. Transition graph of different ecDNA subpopulations**. Blue represents cells carrying one ecDNA type, i.e. pure; purple represents cells carrying both yellow and red elements, i.e. mixed; grey represents cells without ecDNA, i.e. ecDNA-free. The diagram shows the mutual dependency for subpopulations, where the black arrows track the relations in case of simple random segregation and the green arrows add the additional relations with the incorporation of type switching. **c. Selection scenarios**. Quantitative description of the different selection scenarios investigated throughout the paper.

The second key evolutionary parameter in our model is selection, represented by reproduction rates of the yellow and red ecDNA variants *s*_*y*_ and *s*_*r*_ in the range [1, +∞) respectively. Cells without ecDNA have a baseline reproduction rate of 1, while mixed cells carrying both ecDNA variants have a fitness equal to the maximum value of *s*_*y*_ and *s*_*r*_ (Figure 1c). This is because here, we are interested in the maintenance of mixed cell states in the absence of co-selection [16]. Our setting then allows for three different selection regimes: neutral (*s*_*y*_ = *s*_*r*_ = 1), identical positive selection (*s*_*y*_ = *s*_*r*_ > 1) and non-identical positive selection (*s*_*y*_ ≠ *s*_*r*_≥ 1, Figure 1c). During each cell division, ecDNAs are copied and segregated randomly and independently into daughter cells, mathematically modeled by a binomial distribution with probability 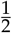 following [8].

### Classification of subpopulations

We define four different subpopulations based on their ecDNA content: pure yellow, pure red, mixed and ecDNA-free cells. A cell is classified as pure if it carries just one ecDNA geno-/pheno-variant, either yellow or red, whilst a cell is defined as mixed if it carries both variants (Figure 1b). At each cell division, a mother cell may generate daughters belonging to different subpopulations, based on the interplay of random ecDNA segregation and switching (Figure 1b). We assume that ecDNA initiation is rare, as we consider complex structures that contain oncogenes and enhancers. A cell losing all ecDNA copies does not gain a new identical copy spontaneously, and its offspring remains ecDNA-free throughout the remainder of the simulation.

## Results

### Single cell EcDNA copy number distributions are sensitive to selection and switching

We first ask how switching rate and selection affect the ecDNA copy number distribution among cells. In a simpler case of only one ecDNA variant, we have previously shown that theory predicts a wide ecDNA copy number distribution consistent with in vitro and in vivo observations of e.g. glioblastoma and neuroblastoma cell line and patient samples [8]. While the shapes of ecDNA copy number distributions keeps being wide when considering multiple ecDNA variants, the concrete distribution is sensitive to selection and switching.

When starting simulations with a single cell containing one yellow ecDNA copy, pure yellow cells dominate the population and have a higher copy number distribution compared to pure red cells in all selection scenarios for small switching probabilities (Figure 2a). We observe a clear difference among neutral, identical and non-identical positive selection, where the copy number distributions of pure cells are highest under non-identical selection, followed by the ones under identical selection and those under neutral selection. The copy number distributions of mixed cells are similar among different selection scenarios probably due to their low abundance. In contrast, for extremely large switching rates (Figure 2c), the copy number distributions of pure yellow and red cells are identical for all selection scenarios, albeit pure yellow cells have higher fitness compared to pure red cells in the case of non-identical positive selection, suggesting that selection plays little role in the dynamics of pure cells when switching rates are extremely high.

**Figure 2.**
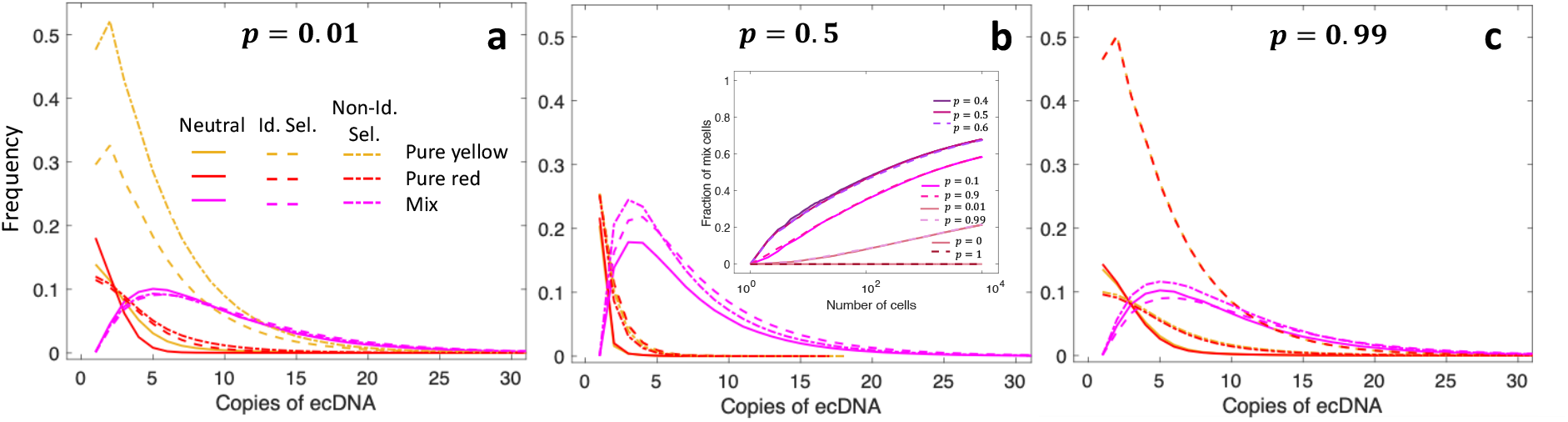
Distribution of ecDNA copies among different subpopulations. We show the ecDNA copy number distributions for pure yellow, pure red and mixed ecDNA cells under small, intermediate and large switching probabilities when *p*_*y*_ = *p*_*r*_ = *p*. For each switching choice, we present results under three selection scenarios defined in Figure 1c with *s* = 2. That is under neutral selection, all cells have fitness as 1; under identical selection, all cells carrying ecDNA have fitness as 2; under non-identical selection, the pure yellow and mixed cells have fitness as 2, while the pure red cells have fitness as 1 as ecDNA-free cells. The internal panel in **b** shows the fraction of mixed ecDNA cells under a broad range of switching probabilities (*p* between 0 and 1). We start our simulations from a single cell with one yellow ecDNA copy, and show the distribution recorded at 10^5^ cells over 1000 realisations.

Somewhat surprisingly, the copy number distribution of mixed cells is quite similar for small and large switching rates (see purple lines in Figure 2). This changes for intermediate switching rates (Figure 2b). Mixed cells (purple lines) dominate for all selection regimes (purple dashed and dashed-dotted lines). The distributions of yellow and red pure cells are narrow and peaked at low copy numbers with very short tails (yellow and red lines).

### The fraction of mixed ecDNA cells is highest under intermediate switching

Intuitively, we would expect that higher switching probabilities will promote higher fractions of mixed ecDNA cells. However, our results show a contrary pattern, where intermediate switching probabilities lead to the highest fraction of mixed cells for all selection regimes (Figure 2b inner panel). We first suspected that this might be related to our initial condition, where the first cell carries only a single copy of yellow ecDNA. Thus, high switching probabilities could lead to synchronous switching of all copies in cells between yellow and red types initially, which dominates the dynamics later.

To exclude the impact of this initial condition, we introduce switching early or late in simulations (Figure 3). When the population grows, there is a fast accumulation of copy number diversity among cells. Switching early or later naturally captures different initial states of copy number distributions where the switching starts to act. Our results show a convergence of the fractions of sub-populations from different switching times under all selection regimes. This convergence is fastest under neutral selection (Figure 3a) followed by identical (Figure 3b) and non-identical positive selection (Figure 3c). This indicates that the fractions of sub-populations approach a composition that is independent of the initial copy number distribution of the population.

**Figure 3.**
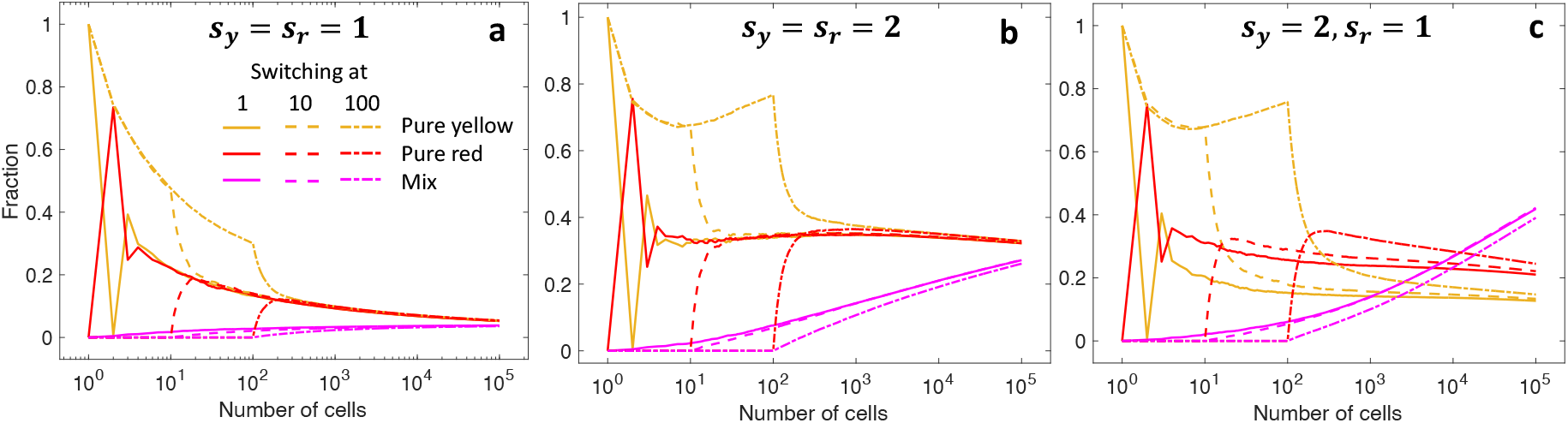
Fraction of subpopulations over time for different starting points of switching. **a**. Neutral selection. **b**. Identical positive selection. **c**. Non-identical selection. In each panel, we compare the fractions of pure and mixed ecDNA subpopulations, when the switching probability is high (*p*_*y*_ = *p*_*r*_ = *p* = 0.99). The start of switching is introduced when the population grows from a single cell to 1, 10 and 100 cells. The initial cell carries one yellow ecDNA copy. All dynamics are averaged over 1000 realisations.

### Maintaining mixed ecDNA cell populations requires both switching and selection

Given the above observations, we now investigate how and when mixed cells can be stably maintained in growing populations. Through the transition diagram between subpopulations (Figure 1b), we see that the ecDNA-free state is an absorbing state under neutral selection with or without switching. This is evident also in Figure 4a, where ecDNA-positive cells rapidly diminish during the population expansion. This is in line with the preliminary discussion earlier and observations in previous work [8, 16]. However, under the same switching probability (*p* = 0.1), and selection for ecDNA-positive cells (Figure 4c), the fraction of mixed cells increases over time.

**Figure 4.**
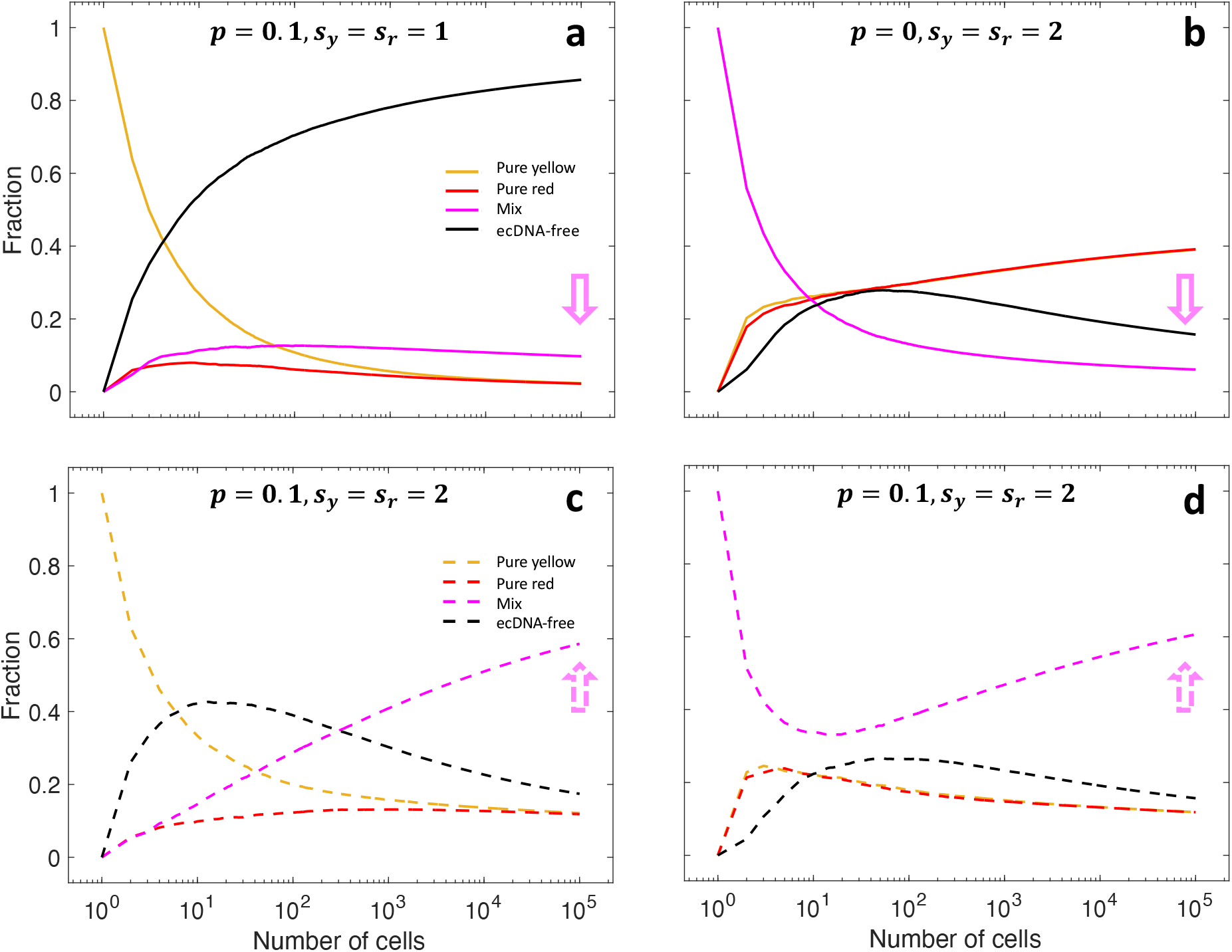
Mixed cells are maintained if selection is positive for cells carrying ecDNA and switching is on. We compare temporal dynamics of the fractions of pure, mixed and ecDNA-free cells when selection and switching are on and off. **a and c**. Starting with a single pure cell, switching is fixed at *p* = 0.1 and selection is neutral in panel a and positive in panel c. **b and d**. Starting with a single mixed cell, selection is positive and fixed at *s* = *s*_*y*_ = *s*_*r*_ = 2 and switching is off in panel b and on in panel d. Following the purple arrows, we can see the fraction of mixed cells only increases if switching and selection are both on, regardless the initial conditions. All dynamics are averaged over 1000 realisations.

Nevertheless, positive selection of cells carrying ecDNA alone is insufficient to maintain a mixed cell population. Figure 4b shows how selection alone still leads to de-mixing in the absence of switching. This corresponds to removing the green arrows in Figure 1b, which makes pure cells an absorbing state under positive identical selection. Only the simultaneous action of positive selection and switching enables the maintenance of the mixed subpopulation (Figure 4c and d).

This observation is further supported by individual realizations rather than averages of the stochastic dynamics. Figure S2 illustrates the fraction of mixed cells given 1000 single realizations of stochastic simulations. Although fluctuations are large, reflecting the stochastic nature of random segregation, all trends qualitatively follow the same pattern: extinction occurs when either switching is 0 or selection is neutral, whereas the populations of mixed cells are consistently maintained when selection is positive and switching is possible.

### Transition between different subpopulations during cell divisions

We then ask what are the probabilities of transitioning between different subpopulations, i.e. pure, mixed, ecDNA-free states, during cell divisions in simulations. We let the population grow to 10^5^ cells and record the frequency of transitions from the state of the mother cell into possible states of both daughter cells (Figures S6-S8) for each division, allowing us to quantify all arrows in Figure 1b. We also record frequencies of daughter cells to remain in a mixed, pure or ecDNA-free state regardless of the nature of their mothers (Figures 5 and S5-S8), thus quantifying the probabilities of staying in each state in Figure 1b.

**Figure 5.**
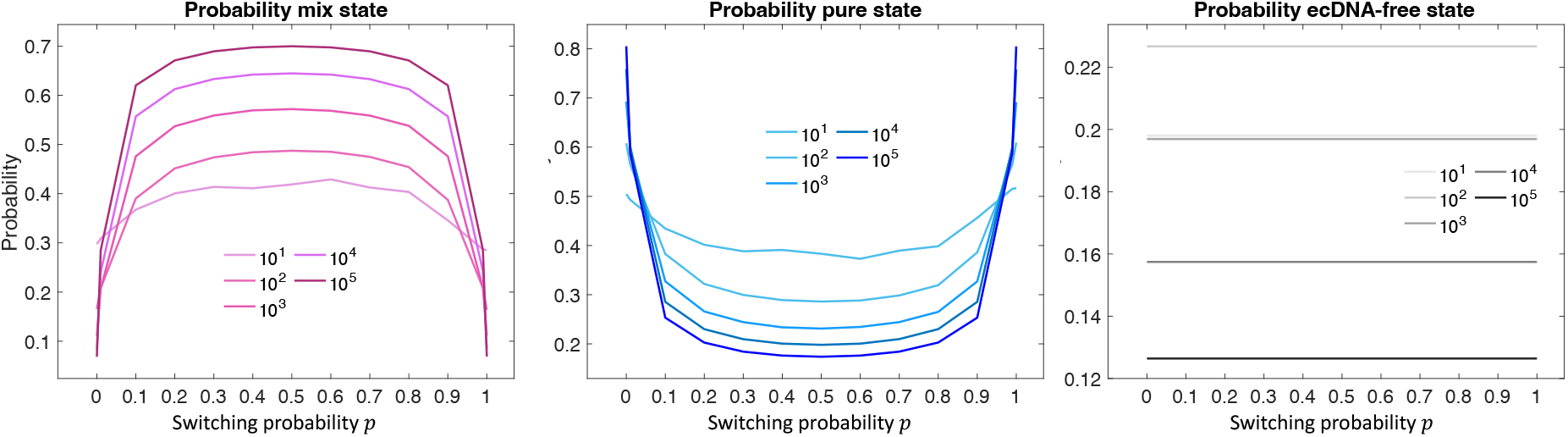
Probability of staying in mixed, pure and ecDNA-free states at different times (population size) under positive identical selection (*p*_*y*_ = *p*_*r*_ = *p*, initial condition: 1 single pure cell with 1 yellow copy). **a**. Probability of staying in mixed state. **b**. Probability of staying in pure state **c**. Probability of staying in ecDNA-free state. These dynamics are generated by simulations starting with a single cell with 1 yellow ecDNA copy and results are recorded over 1000 realisations.

Considering identical positive fitness for both ecDNA types (*s*_*y*_ = *s*_*r*_ = *s*), the probability of being ecDNA-free is independent of the switching probability as expected (Figures 5c and S5c). During the population expansion, the mean ecDNA copy number increases when selection for cells carrying ecDNA is positive. The increasing mean copy agrees with the expected observation that the probabilities of losing ecDNA during cell divisions and thus staying in the ecDNA-free state decrease with increasing population size (Figures 5c and S5c). On the contrary, the probability of staying in the mixed ecDNA state increases over time and peaks for intermediate *p* (Figures 5a and S5a and Figure 2b inner panel). Correspondingly, we observe the opposite behavior for pure ecDNA cells (Figures 5b and S5b). The probability of staying in the pure state (either pure yellow or pure blue cells) has the lowest value under intermediate switching probabilities.

Interestingly, for low (between 0 and 0.05) or high (between 0.95 and 1) switching rates, the probabilities of remaining in a pure state (Figures 5b and S5b) are of opposite order compared to intermediate switching scenarios. We see that in two limiting cases, i.e. ecDNA elements never (*p* = 0) or always (*p* = 1) switch, without additional fitness advantages (co-selection) mixed cells will be lost after the first few divisions and the probability of being in pure states approaches 1. The dynamics of these two extreme cases must be symmetrical, which is confirmed by our simulations starting with different ecDNA configurations in the initial cell (Figure S5).

For intermediate switching rates, transitions from pure into mixed cells are more likely than from mixed into pure cells (Figure S6 d-f), explaining the higher prevalence of mixed cells in that parameter regime. If we vary selection strength (*s* = 2 compared to *s* = 3.5), the probabilities of staying in different sub-populations states change as expected, see Figure S6 black (*s* = 2) and red numbers (*s* = 3.5) inside the circles. A higher selective advantage leads to higher probabilities of staying in ecDNA-positive states (pure and mixed) and a lower probability of being ecDNA-free. However, the selection strength has only a minor impact on the transition probabilities between states.

For neutral selection (*s* = 1), the probability of staying in the mixed state decreases considerably (Figure S7), and mixed cells will get lost over time. Again, the transition probabilities (arrows in the transition diagrams) only deviate marginally from the identical positive selection regime (Figure S6). This further confirms that, without additional fitness advantage (co-selection) for mixed cells, random segregation (black arrows) and switching (green arrows) are the main forces that dominate the transition between different sub-populations rather than selection strength. However, selection drives the reproduction difference between ecDNA-positive and ecDNA-free cells, and thus is also required for the maintenance of mixed cells. Without selection, pure cells upon which switching acts are less abundant, and there is no source to compensate the constant loss of mixed cells due to random segregation.

All these observations remain qualitatively true if we consider different initial conditions, e.g. when the population starts from a single mixed cell of one yellow and one red copy. The highest percentage of mixed cells is still observed under intermediate *p*, with a small quantitative change of transition probabilities and probabilities of staying in different subpopulations (Figure S8).

### Shannon diversity at single-cell level suggests convergence to a homogeneous mixed state

We have studied the composition and copy number distributions of sub-populations and demonstrated how they are impacted by selection and switching. We further explore the diversity in the population by using the Shannon indexes defined at the whole population and single-cell level given by

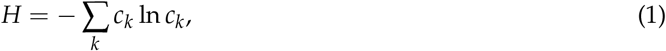

where *c*_*k*_ represents the density of category *k* in the population (Table 1).

**Table 1:** Categories in Shannon index. For the Shannon index at the whole population level, every configuration of copies classified by i yellow and j red copies, is a category, and the number of categories is dynamical ranging from 1 to +∞. For the Shannon index at single-cell level, there are two categories, red and yellow elements.

| Shannon index | Number of categories | Categories |
| --- | --- | --- |
| whole population | dynamical, $[1, +\infty]$ | $\rho_{i,j}$: densities of cells with $i$ yellow and $j$ red ecDNA copies |
| single-cell | 2 | density of yellow ecDNA copies and density of red ecDNA copies in each single cell |

We first look at the Shannon index at the whole population level for different switching probabilities under neutral, identical and non-identical positive selection (Figure S3). We see a clear difference between neutral and positive selection as expected [8]. Under neutral selection, we observe a cluster of low mean ecDNA copy number per cell and small Shannon index values (Figure S3 light grey circles). When selection is positive, the mean ecDNA copy number per cell increases with the fraction of ecDNA-positive cells over time (Figures 3 and 4), reflected by an increasing Shannon index (Figure S3 dark grey and black circles). On the contrary, we hardly observe any distinguishable difference in the Shannon index values across different switching probabilities for all selection scenarios. However, in our analysis above, both selection and switching are required for maintaining mixed cells, and can impact the ecDNA copy number distribution of subpopulations. We conclude that the impact of switching is not reflected in the Shannon index at the whole population level.

We then move to the Shannon index at a single-cell level and look at the diversity of the yellow and red ecDNA copies within mixed cells. Starting from a mixed cell carrying one yellow and one red ecDNA copy, we record the Shannon index values of all mixed cells over time and present their distributions at a given population size. Showing the single-cell Shannon index over its corresponding ecDNA copy number, we have a discrete distribution of all possible values of Shannon index for any given copy number (Figure 6 grey dots). For example, for a mixed cell with 2 ecDNA copies, there is only one possible Shannon index

**Figure 6.**
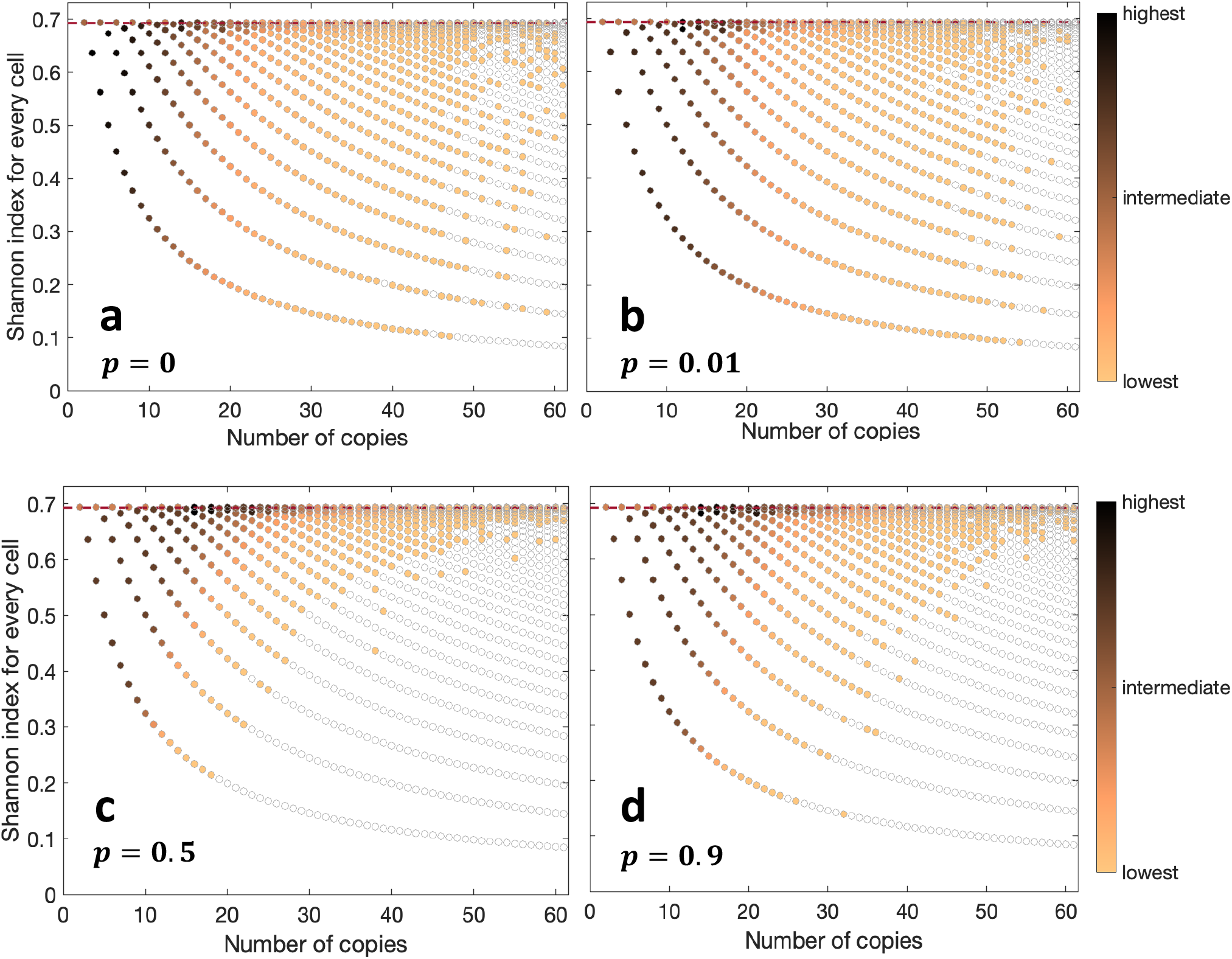
Shannon index at single-cell level. Scatter plots of Shannon index values for four different values of *p* (*s*_*y*_ = *s*_*r*_ = *s* = 2). Each value of the Shannon index is coloured based on its frequency in the population. The horizontal line corresponds to the most frequent Shannon index value. The grey dots on the background are all the possible values on the Shannon index, regardless of *p*. No switching (**a**) versus low (**b**), intermediate (**c**) and high (**d**) switching scenarios are compared. The initial configuration for these simulations is a single cell with 1 yellow and 1 red copy. The indices are recorded at a population size of 10^4^ cells over 600 realisations.

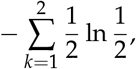

which is around 0.695. This is also the highest Shannon index value a single cell can have independent of its copy number, because the Shannon index decreases when the ratio of yellow and red cells is uneven (Figure 6 horizontal line and Figure S4). In Figure 6, if we join the lowest values of possible Shannon index across all copy numbers, this discrete curve represents all mixed cells with the lowest diversity by having only one yellow or red copy. While their Shannon index will be

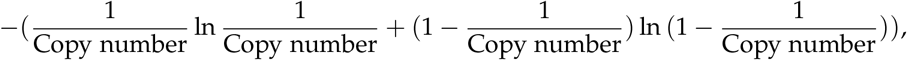

we see a decreasing curve over copy number. A similar logic applies to other discrete curves composed of the grey dots, which refer to the cells with the 2nd, 3rd lowest diversity and so on. Such mixed cells with a low Shannon diversity are more likely to generate pure daughter cells in random segregation, due to the imbalance in copy numbers between the yellow and red ecDNA variants in those cells.

We show the occupation of those possible Shannon index values in stochastic simulations from no switching to high switching probability (Figure 6 coloured dots). Different from the Shannon index at the whole population level, the overall configuration of single-cell Shannon index is significantly influenced by switching. The occupation of low Shannon index curves decreases when the switching probability *p* increases from small to intermediate values (Figure 6 a to b), but increases again when *p* becomes very large (Figure 6 c to d). This demonstrates that the dynamics of mixed cells with low Shannon index values indeed contribute to the transition between mixed and pure cells. Furthermore, we observe an invariant peak in the histogram of Shannon index at its highest possible value (Figure 6 and S4), corresponding to configurations with equal numbers of yellow and red copies in those cells.

This suggests a consistent proliferation behaviour that tends to stabilize toward a more homogeneous configuration between the two ecDNA variants in mixed cells, regardless of the values of the switching probability.

## Discussion

We have built a general framework to model the stochastic dynamics of ecDNA variants in the presence of geno-/pheno-type switching and explore the mechanisms necessary for maintaining mixed cell states in growing populations. Simulations suggest that the coexistence of ecDNA variants within single cells is only possible in the long run if ecDNA positive cells have a selective advantage and can switch (Figure 4). The absence of either condition leads to the extinction of mixed cell states ultimately. As discussed before, there is a second scenario where co-selection allows multi-species ecDNAs to coexist in single cells. A phenomenon observable in about 30% of all ecDNA driven cancers [16]. However, different ecDNA species emerge from different regions of the human genome, excluding the possibility of any switching dynamics to play a role in these circumstances.

Nevertheless, the question of maintenance of ecDNA variants in human cancers in the absence of coselection is of biological importance. There is increasing evidence that point mutations that induce resistance to targeted therapies occur on ecDNAs [6, 32, 33]. It is an open question whether these ecDNA amplifications are pre-existing and selected on or emerge upon treatment. In either scenario, our modeling suggest that upon emergence, mixed cell population of ecDNA variants can coexist. Even worse, given the wide ecDNA copy number heterogeneity, ecDNA presence would immensely increase the likelihood for multiple independent resistant variants to accumulate within the same cell, possibly complicating strategies of targeted combination therapies to overcome resistance evolution [34].

Moreover, about 50% of glioblastoma patients present with *EGFR* amplified on ecDNA [1]. In a high fraction of those, high frequency point mutations and/or deletions are detectable on *EGFR*. These variants tend to be under positive selection and likely are acquired prior to tumour initiation [30], suggesting ecDNA amplifies and probably diversifies in premalignent brain tissue. Similar dynamics have been observed in breast and esophageal cancer [35, 36]. Ultimately, this implies a multi-step process of ecDNA driven tumour initiation in a subset of patients, where first ecDNA forms and then one or a few additional activating events promote tumour initiation [6]. Currently, we neither know details of the premalignent stages in most cases, nor do we know the nature of these additional activating events, with noticeable exceptions in GBM for example. It is however very natural to assume that the interplay of selection and switching and the resulting tendency to maintain mixed ecDNA cell compositions contributes and probably facilitates tumour promotion. We therefore suggest that a better understanding of ecDNA dynamics, details of its accumulation and diversification will also inform our understanding of cancer evolution pre and post tumour expansions.

## Acknowledgements

E.S. acknowledges support by the Engineering and Physical Sciences Research Council (EPSRC) PhD fellowship that is part of the United Kingdom Research and Innovation association (UKRI EP/V520007/1).

B.W. is supported by a Barts Charity Lectureship (grant no. MGU045) and a UKRI Future Leaders Fellowship (grant no.MR/V02342X/1). W.H.is funded by NSFC General Program (grant no. 3217024). This work was delivered as part of the eDyNAmiC team supported by the Cancer Grand Challenges partnership funded by Cancer Research UK (P.S.M. CGCATF-2021*\*100012, B.W. and W.H. CGCSDF-2021*\*100020) and the National Cancer Institute (P.S.M. OT2CA278688, B.W. and W.H. OT2CA278670).

## Code availability

A MATLAB R2024 implementation of the proliferation dynamics of ecDNA variants in a growing population is provided at https://github.com/elisascanu/ecDNA_variants_maintenance.

## Supplementary

### Stochastic simulations

The simulations presented in this paper were conducted using MATLAB (version 2023). Our approach is based on the Gillespie Algorithm [37–40], where cells are selected for proliferation randomly but with a probability proportional to their fitness. More precisely, cells without ecDNA have a baseline proliferation rate of *s* = 1. The fitness effect of ecDNA is incorporated through a proliferation rate *s* > 1 for cells with a selective advantage, 0 < *s* < 1 for those at a disadvantage, and *s* = 1 for no fitness effect. The Gillespie algorithm determines which cell divides and tracks the passage of time, which is then measured in generations. Specifically, cells are grouped based on fitness, and division times are drawn from exponential distributions with parameters determined by selection. The cell with the shortest division time undergoes mitosis, after which time is updated. Thus, each time step corresponds to a division event. Upon selection for proliferation, cells carrying ecDNA double their ecDNA copies, which are then randomly distributed between daughter cells in a binomial trial with a success probability of 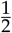. Cells without ecDNA do not pass ecDNA to their daughters, ensuring that ecDNA cannot arise spontaneously in this model. Following the binomial allocation, we consider for each ecDNA copy a type-dependent probability of switching, denoted as *p*_*y*_ for yellow copies and *p*_*r*_ for red copies. For each daughter cell, a random number *r* is drawn for each ecDNA copy; if *r* is lower than the switching probability, the copy changes color. Simulations terminate upon reaching a predefined population size, typically 10^4^ or 10^5^ cells. The recorded output includes the ecDNA copy number per cell at the end of each simulation, allowing for the computation of various analytical metrics such as ecDNA copy distribution, deterministic and stochastic dynamics, and diversity indices.

### Entropy and evenness of the system through Shannon diversity indices

The Shannon diversity index, often referred to simply as Shannon entropy, represents a fundamental measure of the entropy within a system. In the context of ecological studies and population dynamics, the Shannon diversity index provides insights into the diversity, evenness, and distribution of different entities within a population [41–44]. Specifically, when applied to the distribution of ecDNA copies among cells, the Shannon diversity index offers a quantitative assessment of the variability and uniformity in ecDNA abundance across the cellular population.

The formula for this index is:

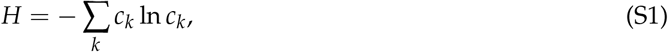

where *c*_*k*_ represents the density of category *k* in the population. By summing the products of each category’s density and the natural logarithm of its density, the Shannon index captures the overall uncertainty or information content associated with the distribution of categories.

It is crucial to note that the Shannon diversity index is sensitive to the relative abundances of different categories within the population. When the abundances of ecDNA types are uneven, with certain variants being more prevalent than others, the resulting Shannon entropy value decreases. Conversely, a more equitable distribution of ecDNA copies among cells leads to a higher Shannon entropy value, indicating greater diversity and evenness in the population.

### Supplementary Figures

**Figure S1:**
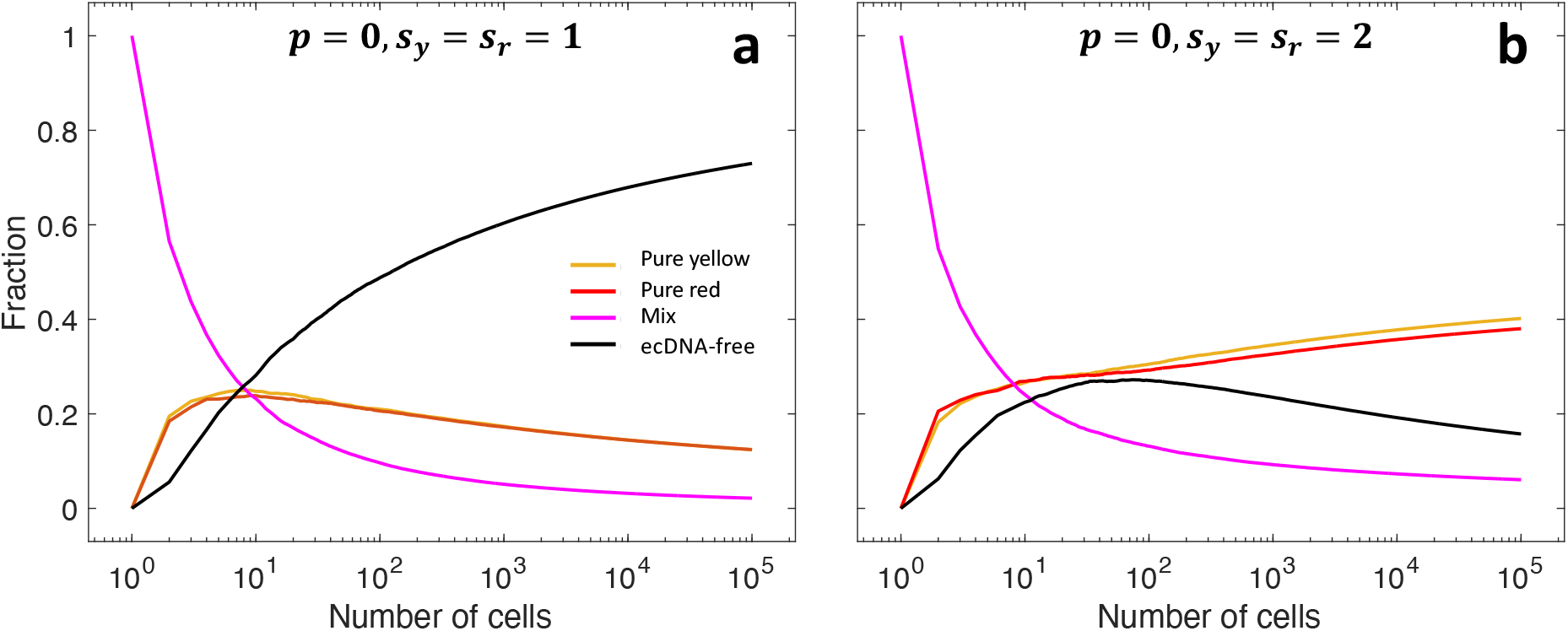
Baseline simulations: de-mixing of the population in absence of switching and further biological assumptions. We analyse temporal dynamics of the fractions of pure, mixed and ecDNA-free cells in the case of multiple ecDNA species, i.e. no switching. **a**. Neutral selection. **b** Positive selection. The initial configuration for both panels is a single mixed cell with 1 yellow and 1 red ecDNA copy. The results are averaged over 1000 realisations.

**Figure S2:**
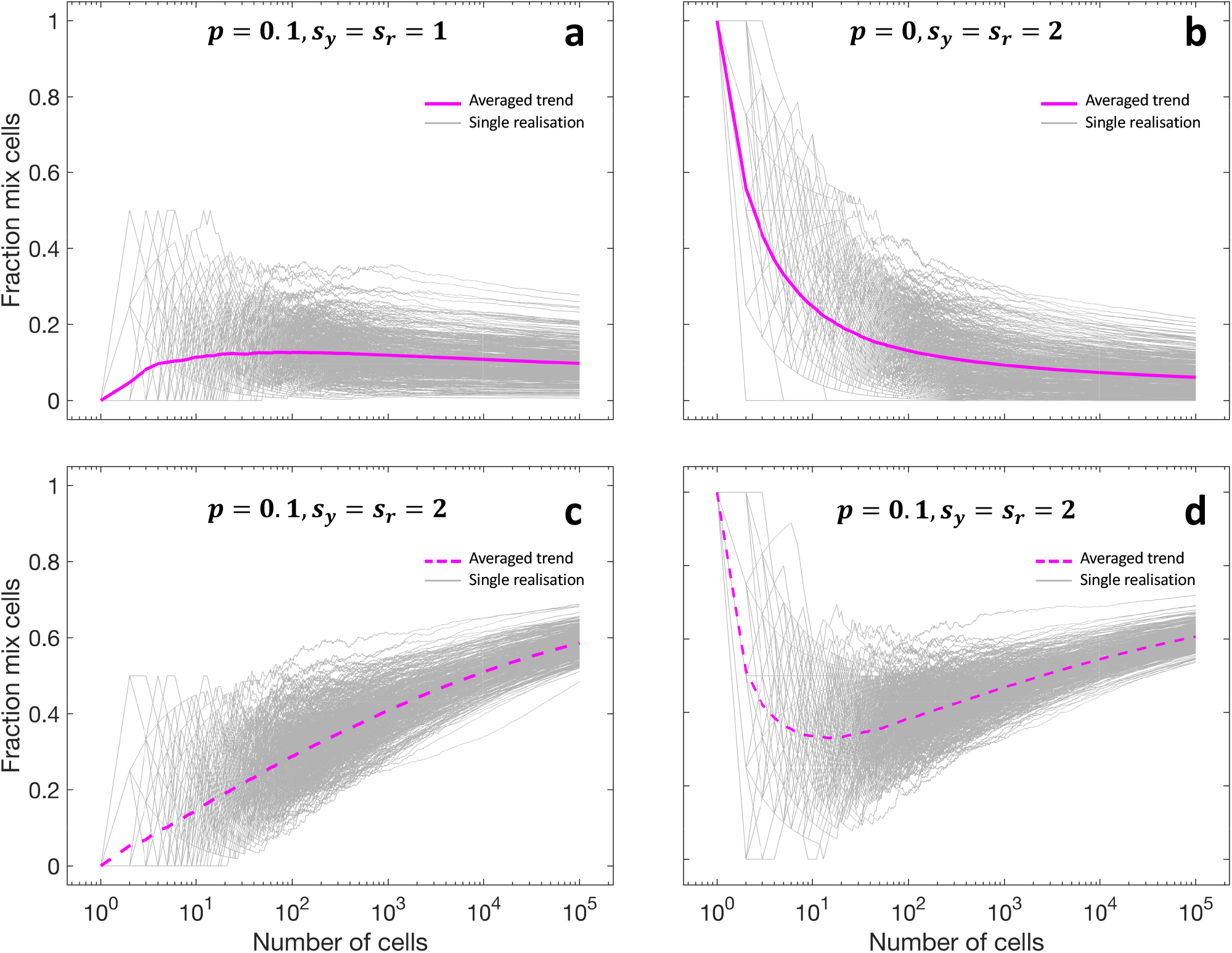
Robustness of claims on multiple ecDNA maintenance in the behaviour of single realisations. We analyse single realisations of growing populations. We record then in light grey the trend of mixed cells for each realisation, and in purple the average among them. The initial condition for simulations in panels **a** and **c** is a single cell with 1 yellow ecDNA copy, whilst in panels **b** and **d** is a single mixed cell with 1 yellow and 1 red ecDNA copy. **a**. Switching is on, selection is neutral. **b**. Switching is off, selection is positive. **c**. Switching is on, selection is positive. **d**. Switching is on, selection is positive. The purple trends are averaged over 1000 realisations.

**Figure S3:**
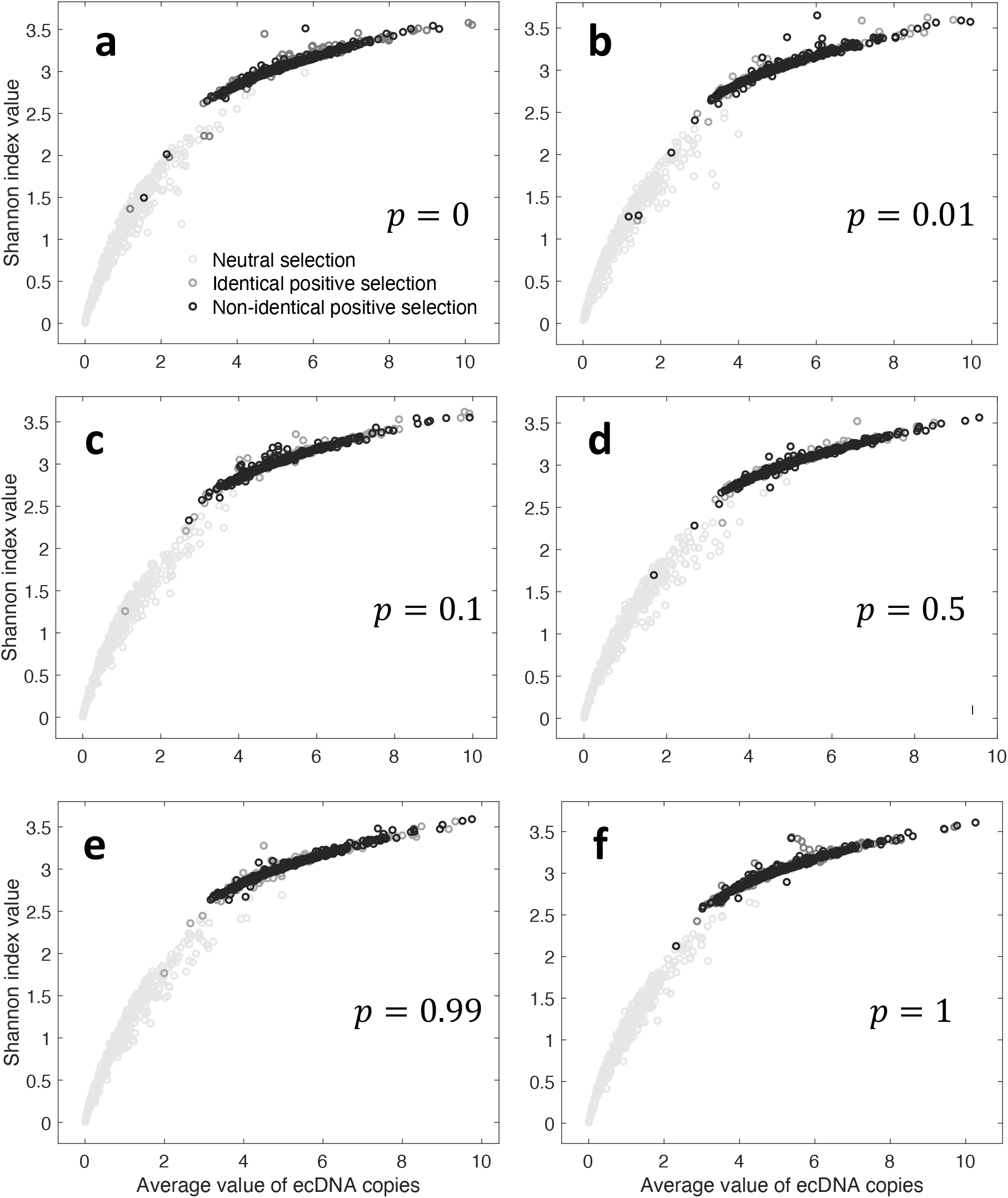
Shannon index at whole population level for different selection scenarios and different values of phenotype switching (*p*_*y*_ = *p*_*r*_ = *p*). Three selection cases are considered: neutral (*s*_*y*_ = *s*_*r*_ = *s* = 1), identical positive selection (*s*_*y*_ = *s*_*r*_ = *s* = 2) and non-identical positive selection (*s*_*y*_ = 2, *s*_*r*_ = 1). Each dot represents the outcome of a single simulation (500 identical repeated realisations per selection scenario in each panel).

**Figure S4:**
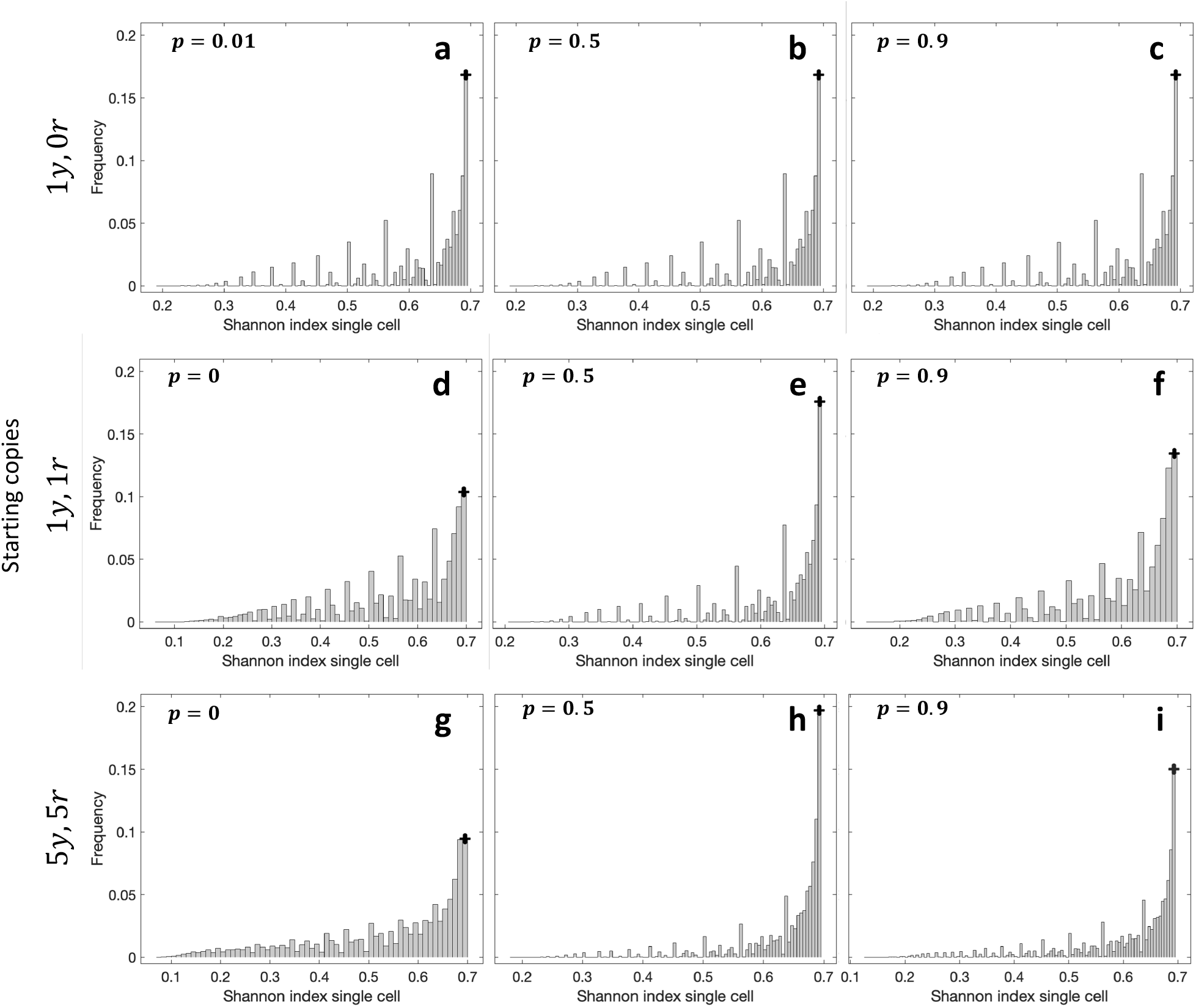
Shannon index at single-cell level. Distribution of Shannon index values for different values of *p* (*s*_*y*_ = *s*_*r*_ = *s* = 2). The red dot on top of a column in each plot identifies the Shannon index value with maximum frequency, which is *H* ≃ 0.635 for every *p*. The initial configuration for these simulations is a single cell with 1 yellow and 1 red copy, and the indices are recorded at a population size of 10^4^ cells over 600 realisations.

**Figure S5:**
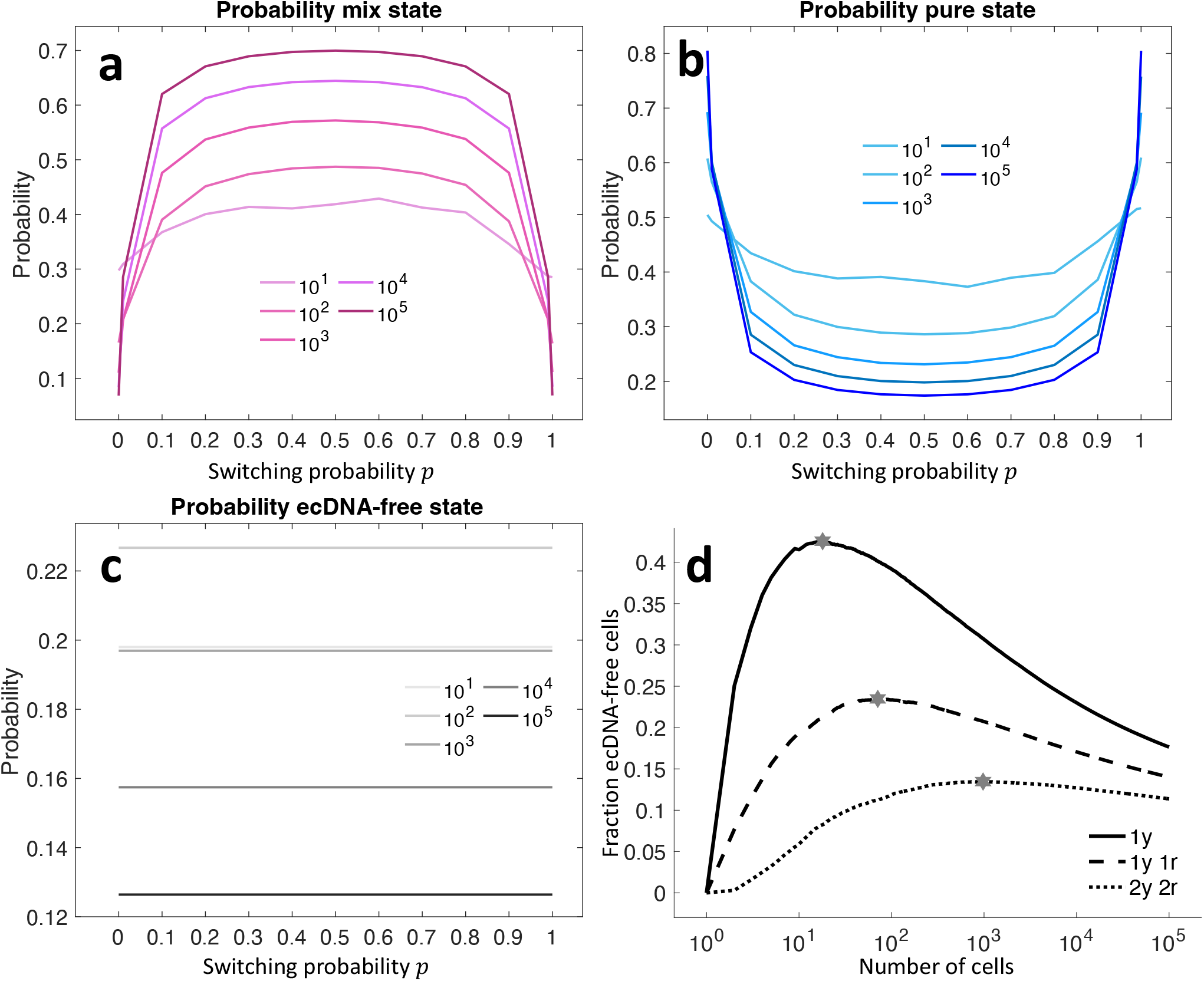
Probability of staying in mixed, pure and ecDNA-free states at different times (population size) under positive identical selection (*p*_*y*_ = *p*_*r*_ = *p*, initial condition: initial condition: 1 single mixed cell with 1 yellow and 1 red copies.). **a**. Probability of staying in mixed state. **b**. Probability of staying in pure state **c**. Probability of staying in ecDNA-free state. **d** Trend of ecDNA-free cells over time for different initial conditions. The star dots indicate the maximum value for ecDNA-free cells density for each curve. These dynamics are generated by simulations starting with a single cell with 1 yellow and 1 red ecDNA copies and results are recorded over 1000 realisations.

**Figure S6:**
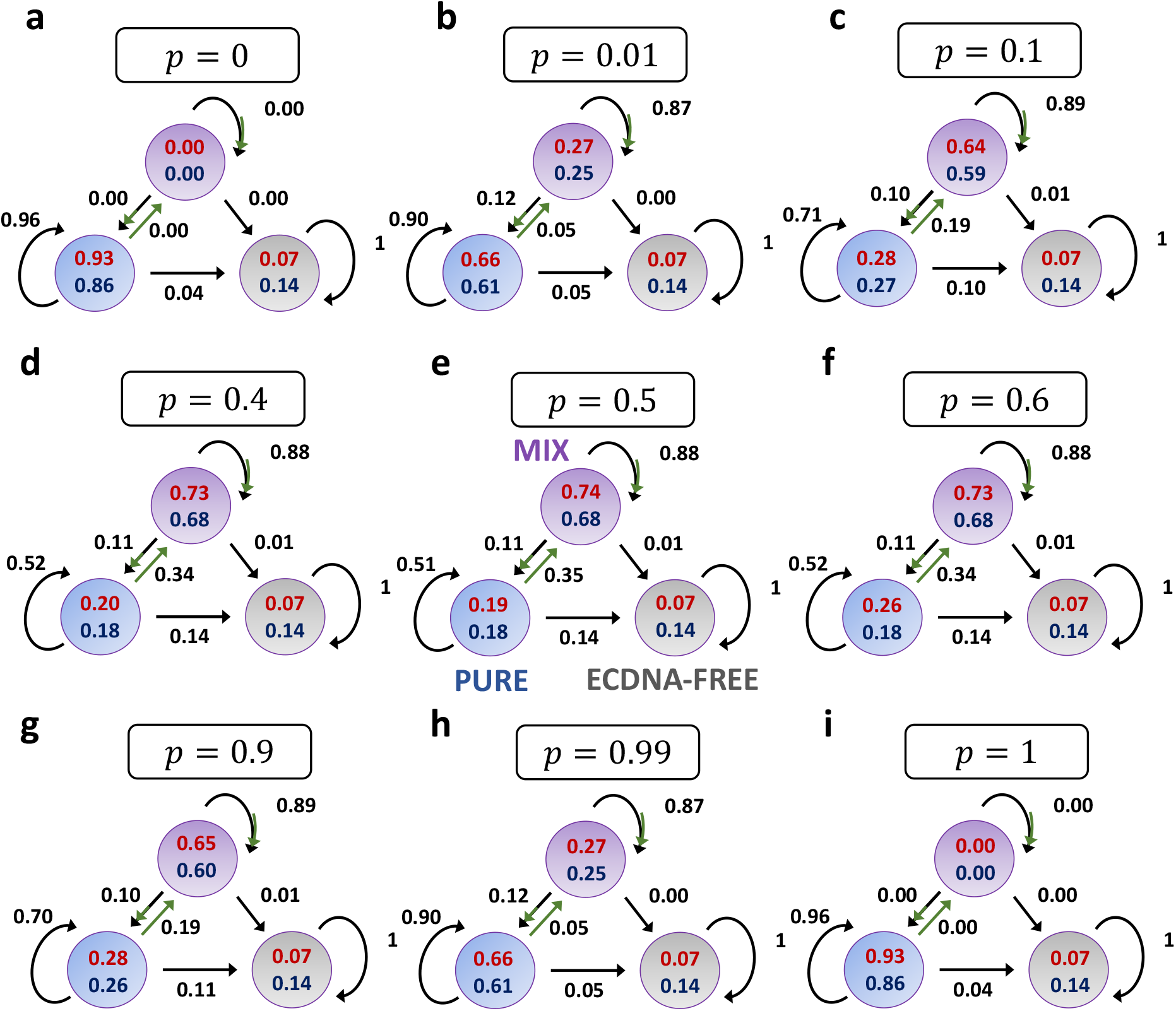
Transition probabilities for model’s dynamics under identical positive selection (*s*_*y*_ = *s*_*r*_ = *s*) recorded at a size of 10^5^ cells (*p*_*y*_ = *p*_*r*_ = *p*, initial condition: 1 single pure cell with 1 yellow copy). The above panels show the probability of transitioning from one state to another, i.e. the arrows’ probability, and the one of staying in a state, i.e. the compartments’ probability, when identical positive selection for both yellow and red ecDNA is considered. Specifically, two selection values are considered, respectively *s* = 2 and *s* = 3.5. Black arrows illustrate the transitions due to random segregation, whilst the green arrows the ones due to switching. The transition probabilities from one state to another, i.e. the arrows, do not change significantly for different positive selection values, whilst the probabilities of staying in a subpopulation, i.e. the compartments, are shown in red colour and blue colour respectively for *s* = 3.5 and *s* = 2. Regardless of the value of selection, when intermediate switching is considered (**d-f**), mixed ecDNA state is the most favoured, with a higher probability of being in its state or transitioning from others, if compared to low (**b-c**) and high (**g-h**) values for *p*. When *p* is at its extreme values, meaning either *p* = 0 (**a**) or *p* = 1 (**i**), then we observe an identical behaviour of the dynamics, with the absence of cells in the mixed ecDNA state and an high transition towards pure state. The initial configuration for these simulations is a single cell with 1 yellow ecDNA copy and results are recorded over 1000 realisations.

**Figure S7:**
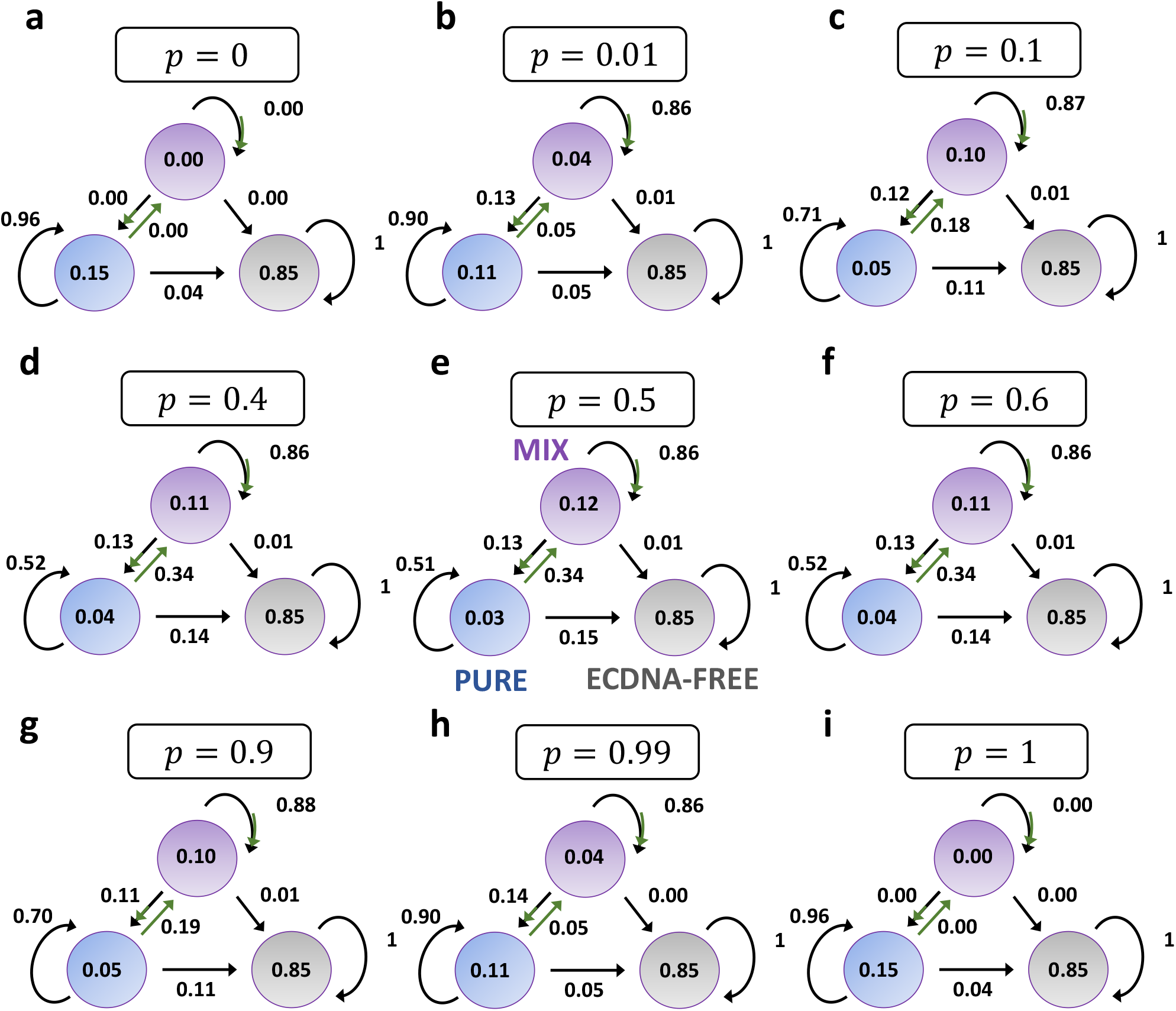
Transition probabilities for model’s dynamics under neutral selection, *s*_*y*_ = *s*_*r*_ = *s* = 1, recorded at a size of 10^5^ cells (*p*_*y*_ = *p*_*r*_ = *p*, initial condition: 1 single pure cell with 1 yellow copy). The above panels show the transition probabilities as in Figure S6 when neutral selection is considered. As in this selection scenario, ecDNA-free cells are the absorbing state, then the transitions and states probabilities change accordingly, compared to positive selection cases. The initial configuration for these simulations is a single cell with 1 yellow ecDNA copy and results are recorded over 1000 realisations.

**Figure S8:**
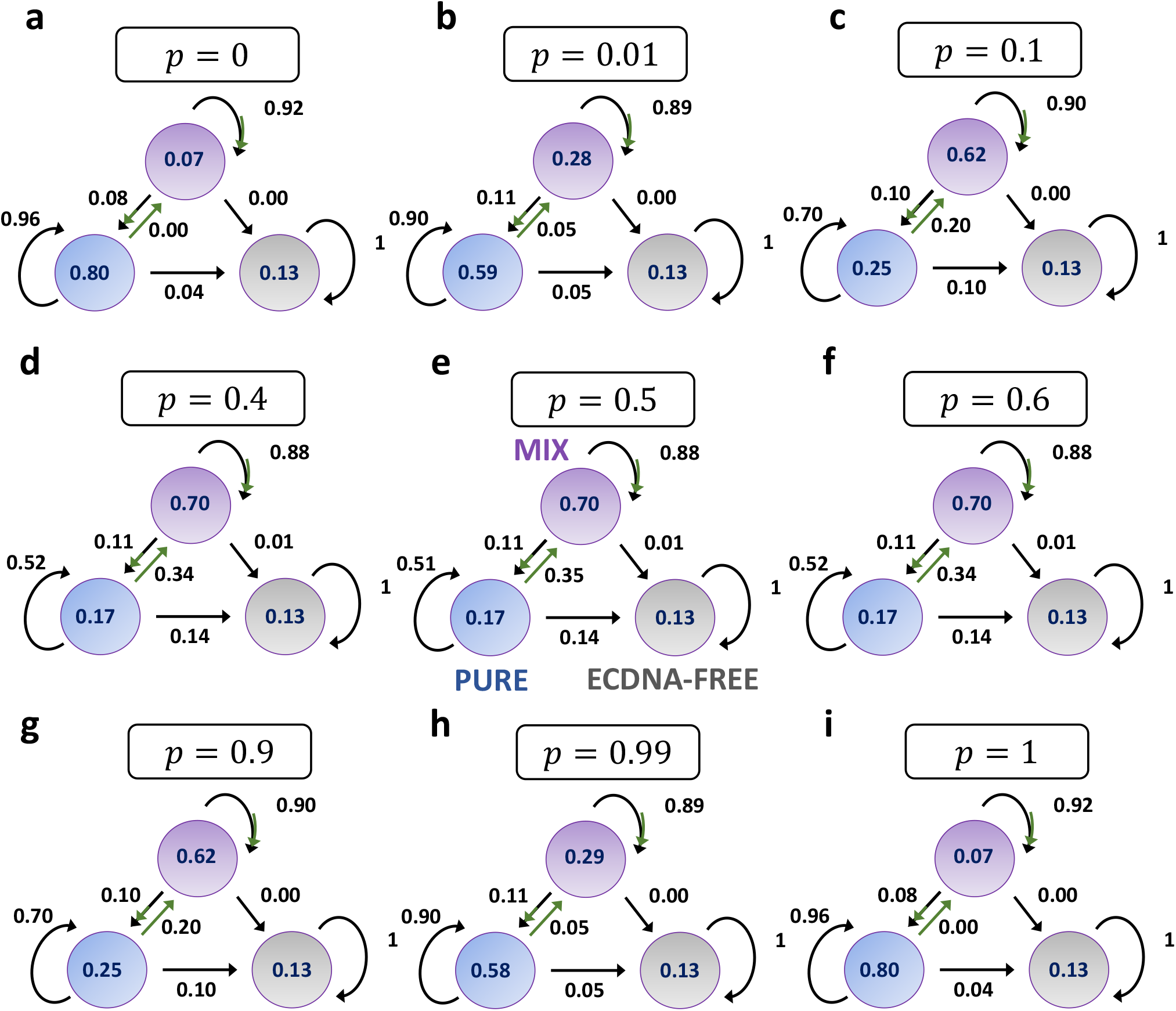
Transition probabilities for model’s dynamics under identical positive selection (*s*_*y*_ = *s*_*r*_ = *s* = 2) recorded at a size of 10^5^ cells (*p*_*y*_ = *p*_*r*_ = *p*, initial condition: 1 single mixed cell with 1 yellow and 1 red copies). The above panels show the transition probabilities as in Figure S6 when another initial condition is considered. We can appreciate the same behaviour in terms of dynamics of mixed cells, as they are more abundant under intermediate switching values (**d-f**). Also, the two extreme cases, i.e. *p* = 0 (**a**) and *p* = 1 (**i**), are symmetric. The initial configuration for these simulations is a single cell with 1 yellow and 1 red ecDNA copies and results are recorded over 1000 realisations.

